# Long-term Humoral Immune Dynamics After SARS-CoV-2 Vaccination and Infection in Healthcare Workers: The VaCoMRI Study

**DOI:** 10.64898/2026.06.10.731348

**Authors:** Mehmet Tekinsoy, Osman Merdan, Rabia Rusen, Catharina Christa, Katharina Müller, Alina Priller, Hrvoje Mijocevic, Hedwig Roggendorf, Otto Zelger, Kathrin Tinnefeld, Samuel D. Jeske, Duluur Vasilev, Sarah Yazici, Johanna Erber, Dieter Hoffmann, Percy A. Knolle, Ulrike Protzer

**Author notes:** **Corresponding Author:** Mehmet Tekinsoy, MD, FEBMM, Institute of Virology, Technical University of Munich, Munich, Germany.

## Abstract

**Objectives:** The VaCoMRI study tracked a healthcare worker cohort from 2020 through June 2024 to characterize the long-term sustainability of vaccine- and infection-induced SARS-CoV-2 immunity.

**Methods:** Following an initial anti-nucleocapsid (N) IgG screening of 4,554 employees in early 2020, participants were monitored regularly from the BNT162b2 rollout (December 2020). Serum collected at predefined intervals after vaccination and breakthrough infection (BTI) was analyzed for anti-N, quantitative anti-spike (anti-S) IgG, and surrogate viral neutralizing antibody (sVNT) titers.

**Results:** Over a median follow-up of 1,180 days, 142 healthcare workers contributed longitudinal samples; 66.2% experienced one BTI and 14.1% a second. In previously seronegative individuals, anti-N titers decreased 2.3-fold between 3 and 6 months post-first BTI, with seropositivity dropping from 82% to 40.4% (median time to seronegativity: 179 days). Conversely, prior anti-N seropositivity was associated with 4.5-fold higher anti-N IgG levels during follow-up, and a second BTI led to 3-fold higher titers at 3 months. Before BTI, the third vaccine dose significantly enhanced antibody persistence versus the second dose; at 9 months, titers remained 7.2-fold higher for sVNT and 3.9-fold higher for anti-S IgG. Initial seropositivity also predicted 2.9-fold higher sVNT levels during follow-up.

**Conclusions:** Repeated vaccination and infection synergistically induced robust, durable spike-directed hybrid immunity. Conversely, rapid anti-N IgG waning severely limits its reliability for the retrospective serosurveillance of SARS-CoV-2 exposure in occupational settings.

## Introduction

Severe Acute Respiratory Syndrome Coronavirus 2 (SARS-CoV-2) emerged in Wuhan, China, in late 2019, spreading worldwide and being declared a pandemic in March 2020. By May 2026, the World Health Organization (WHO) had recorded over 7.1 million reported Coronavirus Disease 2019 (COVID-19) deaths globally (1). In Germany, both the country’s first confirmed COVID-19 case and its first detected Omicron cases were reported in the Munich/Bavaria region (2, 3). The introduction of mRNA vaccines in late 2020 reduced the population risk of severe disease and hospitalization, transforming pandemic control in healthcare and community settings (4). Widespread vaccination campaigns and recurrent waves of infection have created complex layers of immunity across populations. While mRNA vaccines effectively trigger antibodies against the viral spike protein, their levels naturally decline, as demonstrated by multiple longitudinal cohorts (5, 6).

SARS-CoV-2 vaccination elicits spike-directed neutralizing responses, which are shaped by prior infection and booster exposures. Longitudinal cohorts have revealed anti-nucleocapsid (N) IgG declines substantially faster than anti-spike (S) IgG, often resulting in seroreversion. In contrast, S-directed binding and neutralizing activity are more durable, particularly following a booster (7).

Multi-year antibody dynamics inform HCW protection and booster planning. Although mRNA vaccines induce robust initial immunity, long-term studies spanning several years of repeated exposures are limited. Additionally, the rapid decay of anti-N IgG complicates retrospective confirmation of prior infections.

To address these issues, we established the VaCoMRI study (SARS-CoV-2 Vaccination– Induced Immune Response in Employees of the TUM University Hospital rechts der Isar, Munich) (8), a longitudinal cohort of hospital employees that built on the pre-vaccination serostatus from the prior serology screening study “SeCoMRI” (Seroconversion to SARS-CoV-2 in Employees of the TUM University Hospital rechts der Isar, Munich) (9). We closely monitored vaccination and breakthrough infection (BTI) events and focused on interpretable serologic endpoints: anti-N IgG (prior infection), surrogate neutralization, and anti-S IgG.

## Methods

### Study design, participants, and follow-up

At the SeCoMRI study in April–May 2020, 4,554 hospital employees were screened for anti-N IgG. Starting in December 2020, SeCoMRI participants were invited to enroll in the longitudinal VaCoMRI study with follow-up visits scheduled after each vaccination and BTI.

We included HCWs who had received at least 3 vaccine doses and contributed repeated samples through June 2022, with additional follow-up visits captured through June 2024, where available. Prior anti-N seropositivity is defined as a positive anti-N IgG measurement at any given time point before the documented BTI.

### Data collection and serological assays

Detailed vaccination and infection histories were recorded at each visit. Serum samples were collected at standardized time windows following each index event (vaccination or BTI).

To evaluate immune responses, we measured anti-N IgG (iFlash, YHLO; positive ≥10 AU/mL) primarily to determine prior infection, anti-S IgG (Architect, Abbott; positive ≥50 AU/mL, upper limit of quantification (ULOQ)=40,000 AU/mL), and surrogate viral neutralizing antibody (sVNT) titers (iFlash, YHLO; positive ≥10 AU/mL, ULOQ=800 AU/mL).

### Ethics

Both study protocols, including participant information, informed consent form, and information material to promote the study, were approved by the local ethics committee of the Technical University of Munich (ethics vote 476/20 (SeCoMRI) and 26/21S-SR (VaCoMRI)) and conducted in accordance with the Declaration of Helsinki; written informed consent was obtained from all participants.

### Statistical analysis

We evaluated longitudinal antibody kinetics and surrogate neutralization dynamics using linear and Bayesian mixed-effects regression models. The models incorporated fixed effects (e.g., age, sex, measurement timepoint, and prior seropositivity) and participant-level random intercepts to account for repeated measurements.

Further details regarding specific sampling schedules, event handling, assay principles, and statistical modeling parameters can be found in the Supplementary Methods.

## Results

### Cohort and follow-up

We analyzed 142 hospital employees, of whom 94 (66.2%) were female. In 2021, the mean age was 42.8 years with a standard deviation (SD) of 12.4 years. Median age was 44 years in females and 36 years in males. Overall, 65/142 (45.7%) participants were <40 years old, 61/142 (43.0%) were 40–59 years old, and 16/142 (11.3%) were ≥60 years old.

Participants contributed 4,102 serological measurements across 1,775 study visits, averaging 12.5 per participant. Figure 1A-C provides an overview of all serological measurements (Anti-N IgG, Anti-S IgG, and sVNT) across the study period. The median follow-up from first contact to last visit was 1,180 days.

**Figure 1:**
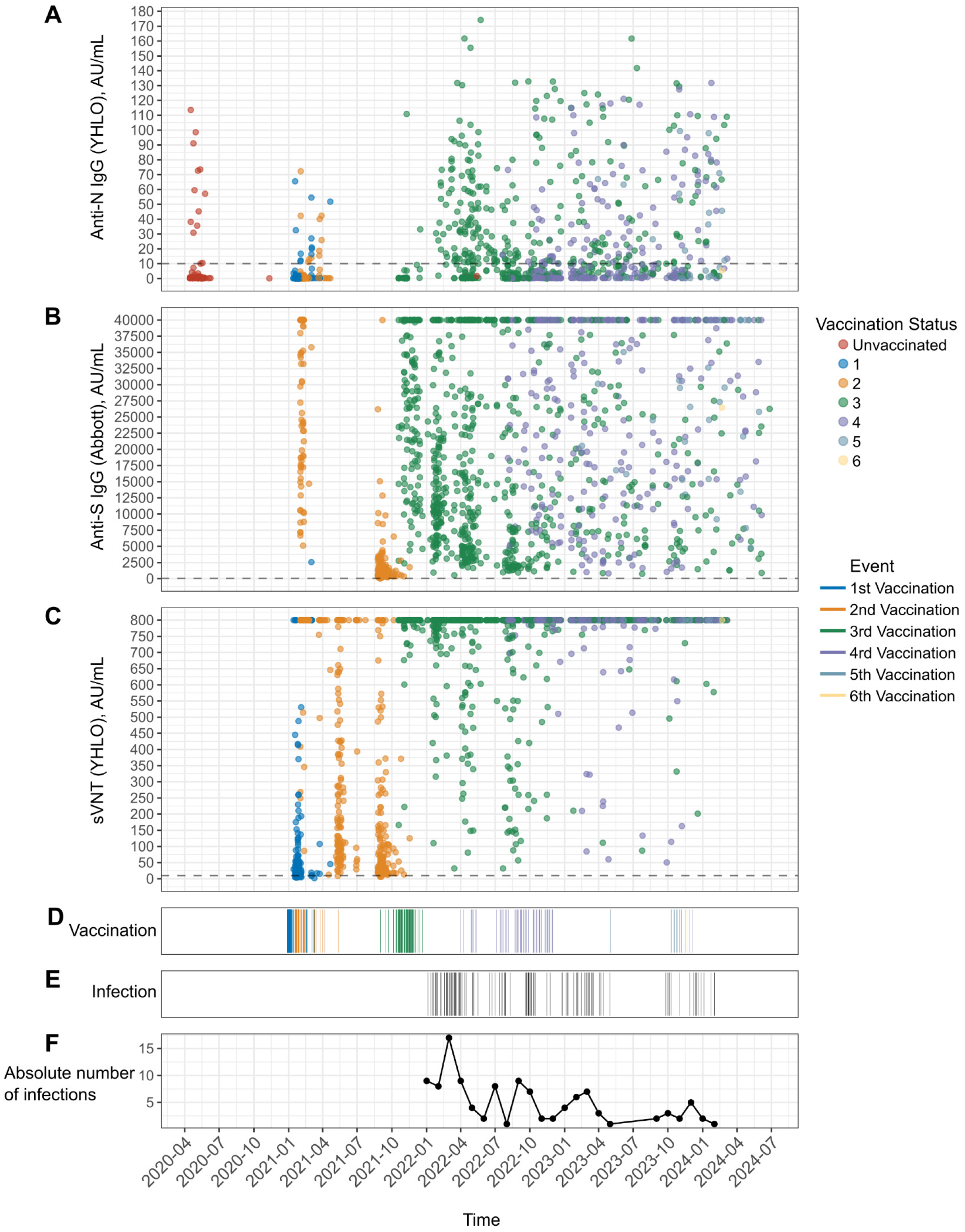
Overview of all data points collected throughout the study. Individual measurements of (A) Anti-N IgG, (B) Anti-S IgG, and (C) surrogate virus neutralization (sVNT) titers are shown, with colors indicating participants’ vaccination status at the time of sampling. Each dot represents an individual measurement. (D) Vaccination and (E) infection events are indicated by vertical lines. (F) Total number of SARS-CoV-2 infections detected per month during the study.

By the end of follow-up, most participants had received three vaccine doses, individual participants up to six vaccine doses: 3 doses in 77/142 (54.2%), 4 doses in 51/142 (35.9%), 5 doses in 13/142 (9.2%), and 6 doses in 1/142 (0.7%) (Figure 1D). The interval between the first and second dose was tightly clustered (median 21 days, IQR 21–22). The median time from the second dose to the third dose was 279 days (IQR: 268–289). Among those who received additional booster vaccinations, the median interval to dose 4 was 321 days (IQR 278–364; n=65) and to dose 5 was 370 days (IQR 354–441; n=14).

### Breakthrough infections (BTIs)

One BTI (1^st^ BTI) was documented in 94/142 (66.2%) participants, and a second BTI in 20/142 (14.1%). Figures 1E and 1F illustrate the temporal distribution of BTIs. At the time of the first BTI, most individuals had received three vaccine doses (74/94, 78.7%). 17/94 (18.1%) participants had received 4 vaccine doses, and 3/94 (3.2%) 5 doses. The time span from the last vaccination to the first BTI was variable, with a median of 144 days (range 28– 683). At second BTI, 14/20 (70.0%) participants had received 3 vaccine doses, and 6/20 (30.0%) had received 4 doses, with a median time since last vaccination of 432 days (range 26–771). The median interval between first and second BTI was 307 days (range 107–696).

Among those who experienced the 1^st^ BTI, 12/94 participants had evidence of prior anti-N seropositivity resulting from exposure to the virus.

### Anti-N IgG Dynamics

In total, 1,225 anti-N IgG measurements were available (Figure 1A). Following the 1st and 2nd BTIs, anti-N titers showed progressive waning with substantial inter-individual variability (Figure 2).

**Figure 2:**
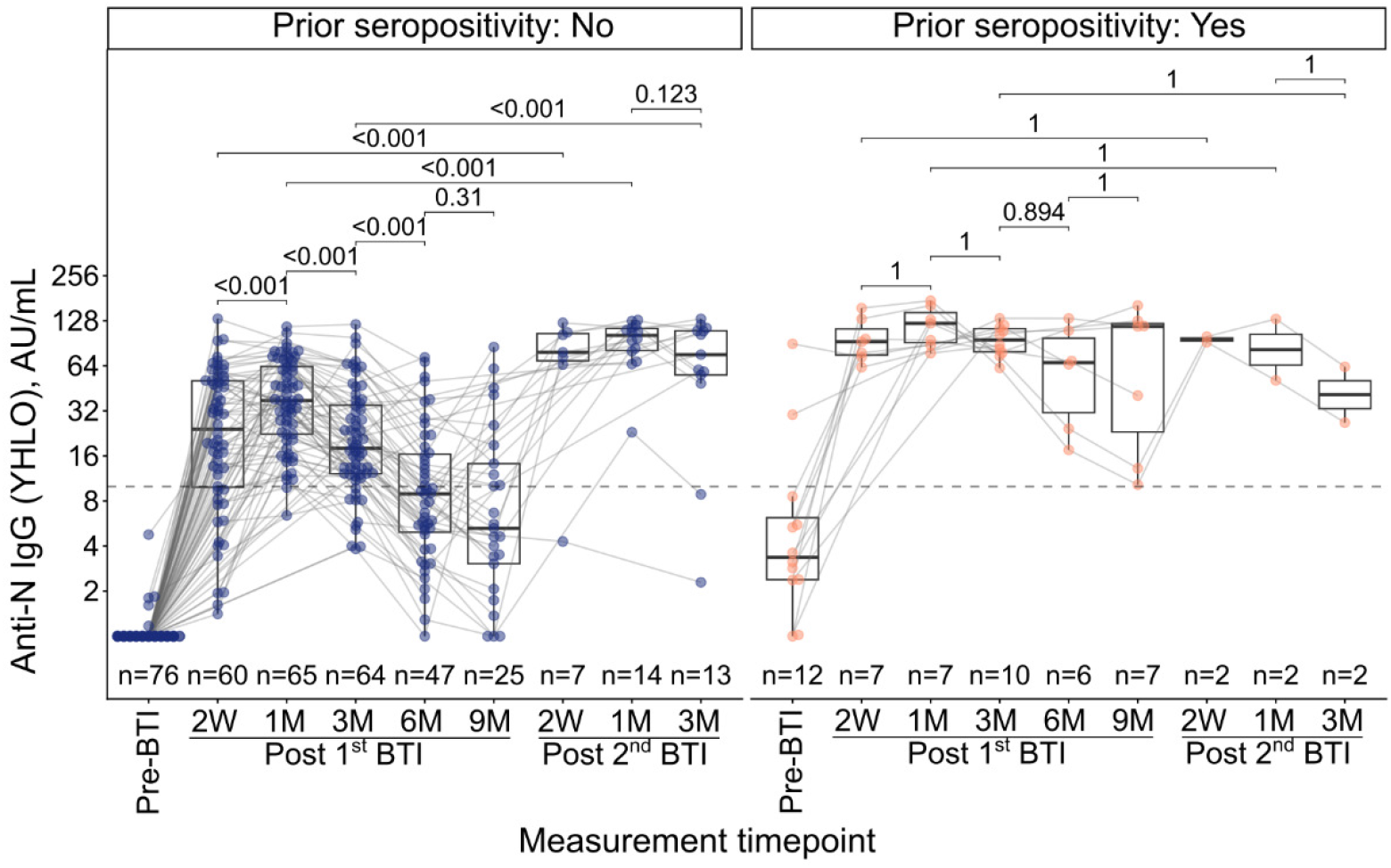
Temporal dynamics of anti-N IgG titers after first and second BTIs. The study group was divided into seropositive and seronegative subgroups based on prior anti-N IgG serostatus, indicating exposure to SARS-CoV-2 before study inclusion. Left panel: no prior anti-N seropositivity, right panel: seropositivity for anti-N. Dots represent individual measurements. Measurements from each subject are connected with lines. Statistical analysis was performed using a linear mixed-effects model with a random intercept for participants. Comparisons between individual time points were made using estimated marginal means derived from the fitted random-intercepts linear model, treating the measurement time point as a categorical variable. Holm-adjusted p-values for the corresponding planned comparisons were displayed. Y-axis values were log 2-scaled. For each measurement time point, the number of available samples (n) is shown. Horizontal dashed lines indicate the threshold value (10 AU/mL) for a positive test result. Pre-BTI marks the most recent available measurement before the detection of the first BTI. AU: Arbitrary units. W: Week(s); M: Month(s), BTI: Breakthrough infection.

Anti-N IgG titers in participants without prior anti-N seropositivity peaked between 2 and 4 weeks and declined between 1 and 6 months, with a significant boosting effect after the 2^nd^ BTI (Figure 2). Age and gender were not significantly associated with anti-N IgG levels (p-values: 0.74 and 0.44, respectively). Individuals with prior anti-N seropositivity had estimated 4.5-fold higher anti-N IgG levels (95% CI:2.5-8.3, p<0.001), showing a booster effect upon repeat exposure.

Among participants without prior infection, model-based estimates indicated a 1.8-fold (95% CI: 1.3-2.4; Holm-adjusted p<0.001) decrease between 1 and 3 months after the 1^st^ BTI, and a 2.3-fold (95% CI: 1.6-3.1; Holm-adjusted p<0.001) decrease between 3 and 6 months. Parallel to titers, the seropositivity rate declined between post-1^st^ BTI 3 months (82%, 53/64) and 6 months (40.4%, 19/47). Overall, the first negative anti-N after the 1^st^ BTI was recorded at a median of 179 days (N=54, IQR=15-239).

One month after the 2^nd^ BTI, titers were estimated to be 2.6-fold (95% CI: 1.5-4.3; Holm-adjusted p<0.001) higher than the same time point after the 1^st^ BTI. Comparison of titers three months after 2^nd^ BTI versus 1^st^ BTI revealed approximately a 3-fold increase (95% CI:1.72-5.12; Holm-adjusted p<0.001).

### Anti-spike IgG

In total, 1,293 quantitative anti-S IgG measurements were collected (Figure 1B). To compare S antibody kinetics after the second and third doses, only measurements obtained before any BTI were included (Figure 3A).

**Figure 3:**
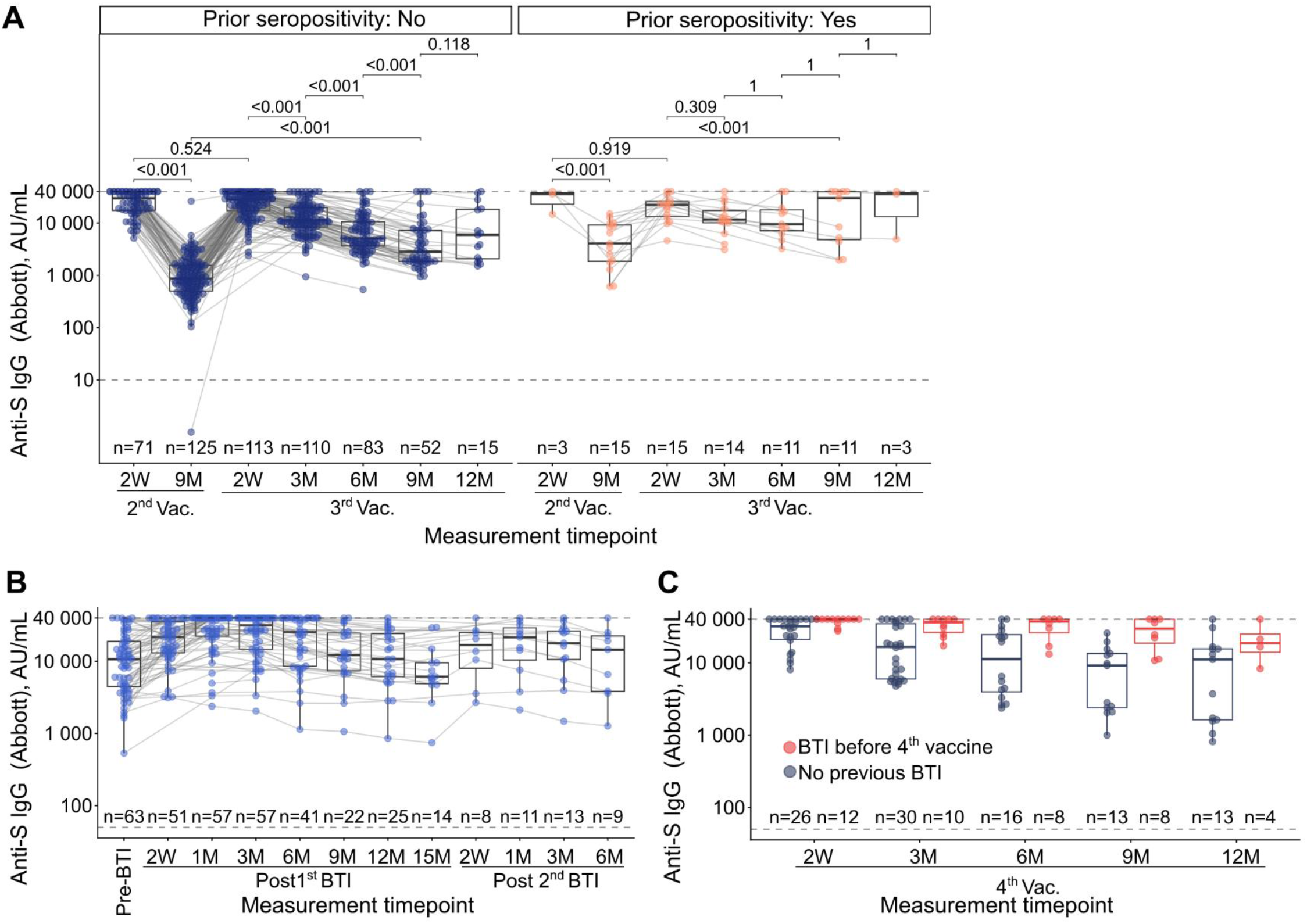
Temporal dynamics of anti-S IgG titers. A. Change in anti-S IgG titers after 2^nd^ and 3^rd^ vaccination dose, factored for prior seropositivity of anti-N IgG representing prior exposure to SARS-CoV-2, was presented. For selected time points, all available measurements before any BTI were shown. Dots represent individual measurements. Measurements from each subject are connected with lines. Statistical analysis was performed using a linear mixed-effects model. Comparisons were made using estimated marginal means derived from the fitted random-intercepts linear model, treating the measurement time point as a categorical variable. Holm-adjusted p-values for the corresponding planned comparisons were displayed. B. Anti-S IgG titer traces of post BTI measurements among participants who received three vaccinations. Pre-BTI measurements are the most recent measurements taken before the BTI, which are no more than 120 days old. C. Summary of the measurements taken after the 4th vaccination before a new BTI. Measurements were colored based on the previous BTI history after the 3^rd^ vaccination. Y-axis values were log 10-scaled. For each measurement time point, the number of available samples (n) was shown. Horizontal dashed lines indicate the threshold value (50 AU/mL) for a positive test result, and the assay upper limit of quantitation (4000 AU/mL). W: Week(s); M: Month(s), Vac: Vaccination

In participants without prior anti-N seropositivity, anti-S IgG titers declined significantly between 2 weeks and 9 months after the second dose, with an estimated 27.5-fold reduction (95% CI: 21.3–34.7; Holm-adjusted p<0.001) (Figure 3A). Because 32.4% (24/74) of measurements at 2 weeks post-second-dose were at the assay ULOQ. Third booster dose restored antibody titers to a similar level 2 weeks after the 2^nd^ vaccination (Holm-adjusted p=0.638) (Figure 3A). Two weeks after the 3^rd^ vaccination, 22% (25/133) of measurements were at the assay ULOQ (anti-S IgG titer IQR: 17,325-37,486). Compared to the 2^nd^ dose, a slower waning of titers was observed after the 3^rd^ dose, consistent with observations of sVNT titers. Post 3^rd^ vaccine dose between 2 weeks and 3 months, 3 to 6 months and 6 to 9 months model-based estimates showed 1.89 fold (95% CI: 1.52-2.35; Holm-adjusted p<0.001), 1.9 fold (95% CI: 1.53-2.46; Holm-adjusted p<0.001) and 1.6 fold (95% CI: 1.2-2.3; Holm-adjusted p<0.001) decrease respectively. Notably, at the 9-month mark, titers remained 4.2-fold higher following the third dose compared to the second dose (95% CI: 3.2-5.6; Holm-adjusted p<0.001) (Figure 3A).

Prior anti-N seropositivity and gender were not statistically significant predictors of antibody titer (p=0.23 and 0.24, respectively). However, prior anti-N seropositivity and measurement timepoint interaction effects were significant at 9 months after the 2^nd^ vaccination, implying almost 6-fold higher titers (p=0.0018). Age was a relevant factor (p=0.0083); a 1 SD (12.4 years) increase in age was associated with a 13% decrease (95% CI: 4%-23 %) in titer.

Post-1st BTI in triple-vaccinated participants, anti-S IgG titers peaked around 1 month, when 50.9% (29/57) of measurements were at the assay ULOQ (IQR: 22,319-40,000) (Figure 3B). After 12–15 months, values approached those before BTI levels.

All measurements after the 3^rd^ vaccination were above the assay-positive threshold. After the 4^th^ vaccination, anti-S IgG measurements were scarce (Figure 3C). Participants were divided into two groups: one with BTI before the 4^th^ vaccination and the other with no prior documented infection. Exploratory analysis showed a trend toward slower antibody titer decline among participants with prior BTI.

### Surrogate viral neutralizing antibodies dynamics

In total, 1,584 sVNT measurements were available after vaccination (Figure 1C). To evaluate longitudinal vaccine-induced immunity before any BTI, we analyzed measurements from infection-free participants at 2 weeks post-1^st^ dose, 3 and 6 months post-2^nd^ dose, and 6 and 9 months post-3^rd^ dose.

Two weeks after the 1^st^ vaccination, the median sVNT titers were 29.2 (n=122, IQR: 16.1– 61.1) and 800 AU/mL (n=16) in participants without and with prior anti-N seropositivity, respectively (Figure 4A). Following the 2^nd^ vaccination, almost all measurements reached the assay ULOQ (Figure 4A), and three months later, the median titer dropped to 188 AU/mL among participants without prior anti-N seropositivity. In this group, a mean 3.5-fold decline (95% CrI:3.1-3.8) was observed between 3 and 9 months; titers at 9 months were 1.7-fold (95% CrI:1.46-1.9) higher than at 2 weeks post-1^st^ vaccination (Figure 4A).

**Figure 4:**
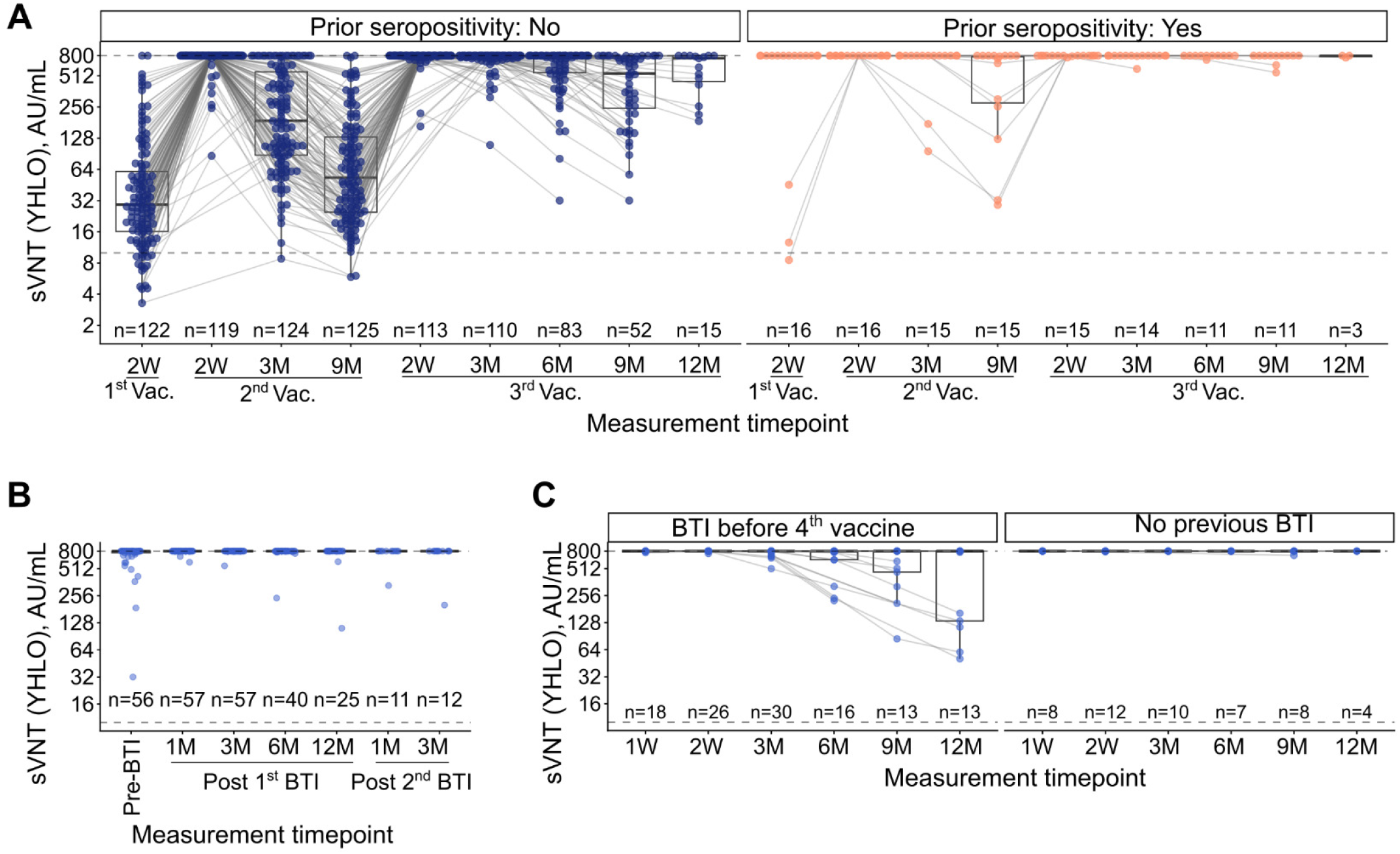
Surrogate virus neutralization (sVNT) titers after vaccination and BTI. A. The study group was divided by prior seropositivity for anti-N IgG, indicating prior exposure to SARS-CoV-2. Left panel: no prior anti-N seropositivity, right panel: seropositivity for anti-N. sVNT titers (YHLO, AU/mL) over time in study participants without documented breakthrough infection were displayed. B. sVNT titers over time among participants with first and second BTI after 3^rd^ vaccination. Pre-BTI measurements are the most recent measurements taken before the BTI, which are no more than 90 days old. C. Summary of the measurements taken after the 4th vaccination before a new BTI. Measurements were facetted based on the previous BTI history after the 3^rd^ vaccination. Y-axis values were log 10-scaled. For each measurement time point, the number of available samples (n) was shown. Horizontal dashed lines indicate the threshold value (10 AU/mL) for a positive test result, and the assay upper limit of quantitation (800 AU/mL). W: Week(s); M: Month(s), Vac: Vaccination

The third dose provided a much larger boost than the second. sVNT titers remained at or very close to assay ULOQ for at least 3 months (Figure 4A). Despite a significant 2.5-fold (95% CrI:1.7-3.7) decay being observed between 6 and 9 months, antibody levels at 9 months after the 3^rd^ vaccine dose were 7.2-fold higher (95% CrI:5.4-9.5) than those recorded after the 2^nd^ dose. Overall, after the 2^nd^ vaccination dose, except for 3 measurements, all analyzed measurements exceeded the assay-positive threshold.

Prior anti-N seropositivity strongly predicted higher responses, yielding 8-fold higher titers (CrI: 3.1-23.5) nine months post-3rd vaccination. A 1-SD increase in age (12.4 years) reduced titers by 39% (CrI: 26%-48%), while gender effects were negligible.

sVNT titers after BTIs post-3^rd^ dose were consistently at or very close to the ULOQ. Six months after 1^st^ BTI, 82.5% of sVNT measurements (33/40) were at ULOQ. Similarly, 12 months after 1^st^ BTI, 88% of sVNT measurements (22/25) were at ULOQ.

Parallel to observations of post-3^rd^ vaccination titers among BTI-free individuals, neutralizing antibody titers remained at or near the assay ULOQ after the 4^th^ vaccination. However, some individuals showed a visible decay of sVNT titers after 3 months. Although measurements were limited for individuals who experienced a BTI between the 3^rd^ and 4^th^ vaccine doses, the available data showed that their titers remained consistently at the ULOQ.

## Discussion

In this longitudinal cohort of 142 hospital employees followed over nearly four years, we observed contrasting longitudinal dynamics between vaccine-induced and infection-induced humoral immunity. Repeated SARS-CoV-2 vaccination elicited robust, highly neutralizing, and sustained spike-directed antibody responses (anti-S IgG and sVNT), with a significantly slower waning rate than after the primary vaccination series. However, these persistent antibody levels did not guarantee sterilizing immunity, as evidenced by the high incidence of BTIs during the follow-up period. Furthermore, unlike the durable spike-directed responses, infection-induced anti-N levels waned rapidly, with a substantial reduction in seropositivity within six months post-exposure.

A key finding of our study is the clear divergence between spike-directed and nucleocapsid-directed antibody trajectories. Following BTI, anti-N increased but then declined rapidly, resulting in a significant drop in seropositivity rates between three and six months after the initial BTI. This pattern is consistent with previous studies showing that anti-N antibodies are less persistent than spike-directed responses and often become undetectable during follow-up (7). In contrast, both anti-S IgG and sVNT remained detectable at substantially higher levels over time, particularly after booster vaccination.

Our findings also highlight the importance of prior SARS-CoV-2 exposure in shaping subsequent humoral responses. Participants with prior anti-N seropositivity had higher anti-N IgG and sVNT levels over follow-up, and a larger proportion remained at the ULOQ for sVNT after the third dose. These results are consistent with the broader literature showing that hybrid immunity is associated with stronger and more sustained immune responses than vaccination alone. In the SIREN cohort, for example, previously infected individuals who were subsequently vaccinated maintained high levels of protection against reinfection for more than one year (10).

Our longitudinal observations of the primary vaccine series reinforce the established global consensus that mRNA vaccination triggers a robust initial humoral response, which predictably wanes over the subsequent 6–12 months (6, 11-15). The third vaccine dose, however, was a particularly important inflection point in our cohort. Compared with the second dose, it was associated with significantly higher anti-S IgG and sVNT responses, with many sVNT measurements reaching or approaching the assay ULOQ. This finding is consistent with published data on the immunogenicity of booster doses and with studies showing enhanced humoral responses after additional vaccine doses (16, 17). Although some measurements reached the assay ceiling, limiting discrimination of the highest responses, the overall pattern strongly supports preserved humoral recall capacity in this cohort.

Despite the strengths of our nearly four-year longitudinal follow-up and the high density of serial samples, several limitations must be acknowledged. First, the single-center design and the cohort composition, which consisted of relatively healthy hospital employees with high vaccine uptake and ongoing occupational exposure, limit generalizability. In addition, the present analysis was based on a subgroup of 142 participants who underwent repeated longitudinal sampling. This may favor individuals with greater follow-up adherence, potentially introducing selection bias. Second, serological measurements were based on commercial assays with inherent constraints. After booster vaccination, a substantial proportion of anti-S and sVNT values reached the assay ULOQ, which limits discrimination at peak response. Although the surrogate neutralization assay provides a practical approximation of neutralizing activity, it does not fully capture the complexity of live-virus neutralization. Third, infection ascertainment was heterogeneous and included institutional PCR, external PCR, and self-reported positive antigen tests. As a result, some asymptomatic or unreported infections may have been missed, and infections were not analyzed separately by diagnostic modality. In addition, because blood sampling occurred at predefined but pragmatic intervals, the precise timing of infection and serologic peak responses could not always be determined. Fourth, our study focused on humoral immunity and did not include measurements of cellular immunity. However, T-cell responses remain relevant, even when antibody levels decline. Finally, the study spanned multiple epidemiological phases, including the emergence of antigenically distinct variants, which may have influenced the apparent waning through both biologic decline and changing antigenic match over time.

In this multi-year cohort of healthcare workers, repeated SARS-CoV-2 vaccination induced strong and relatively sustained spike-directed humoral responses, particularly after the third dose, whereas infection-associated anti-N IgG declined substantially faster after BTI. Prior SARS-CoV-2 exposure was associated with stronger subsequent humoral responses, consistent with a hybrid immunity pattern. Ultimately, although anti-N IgG is useful for identifying recent infections, establishing multi-year or decadal serological footprints requires further research. Additionally, since sustained spike-directed responses do not reflect sterilizing immunity despite their high prevalence, future research should focus on developing alternative serological markers to comprehensively evaluate long-term protection.

## Data availability

The study protocol for VaCoMRI is available from the authors upon reasonable request. De-identified data supporting the findings of this study can be obtained from the corresponding author upon reasonable request and subject to institutional approvals.

## Acknowledgments

We thank all hospital employees who participated, the NoCoV outpatient clinic team, and the diagnostics laboratories for assay support.

## Author contributions

Conceptualization: P.A.K.; U.P.

Methodology: M.T.; C.C.; K.M.; A.P.; H.M.; P.A.K.; U.P.

Investigation: M.T.; O.M.; R.R.; C.C.; K.M.; A.P.; S.D.J.; H.M.; J.E.; H.R.; O.Z.; K.T.; D.V.; S.Y.; D.H.

Data curation: M.T.

Project administration: U.P.; P.A.K; C.C.; M.T.

Formal analysis: O.M.; M.T.

Visualization: O.M.

Writing-original draft: M.T., O.M.

Writing-review & editing: all authors

Supervision: U.P., P.A.K.

## Funding

Supported by the participating institutes and the German Center for Infection Research (DZIF).

## Conflicts of interest

The authors declare no competing interests.

## References

1. World Health Organization. WHO Coronavirus (COVID-19) Dashboard: Deaths. 2026 [updated 2026 May; cited 2026 May 26]. Available from: https://data.who.int/dashboards/covid19/deaths?n=o.

2. Koch-Institut BLfGuLR. Beschreibung des bisherigen Ausbruchsgeschehens mit dem neuartigen Coronavirus SARS-CoV-2 in Deutschland (Stand: 12. Februar 2020). [Description of the initial outbreak of the novel coronavirus SARS-CoV-2 in Germany (as of 12 February 2020)]. Epidemiologisches Bulletin. 2020.

3. Dachert C, Muenchhoff M, Graf A, Autenrieth H, Bender S, Mairhofer H, et al. Rapid and sensitive identification of omicron by variant-specific PCR and nanopore sequencing: paradigm for diagnostics of emerging SARS-CoV-2 variants. Med Microbiol Immunol. 2022;211(1):71–7.

4. Lin DY, Gu Y, Wheeler B, Young H, Holloway S, Sunny SK, et al. Effectiveness of Covid-19 Vaccines over a 9-Month Period in North Carolina. N Engl J Med. 2022;386(10):933–41.

5. Choi MJ, Heo JY, Seo YB, Yoon YK, Sohn JW, Noh JY, et al. Six-month longitudinal immune kinetics after mRNA-1273 vaccination: Correlation of peak antibody response with long-term, cross-reactive immunity. Front Immunol. 2022;13:1035441.

6. Srivastava K, Carreno JM, Gleason C, Monahan B, Singh G, Abbad A, et al. SARS-CoV-2-infection- and vaccine-induced antibody responses are long lasting with an initial waning phase followed by a stabilization phase. Immunity. 2024;57(3):587–99 e4.

7. Van Elslande J, Oyaert M, Lorent N, Vande Weygaerde Y, Van Pottelbergh G, Godderis L, et al. Lower persistence of anti-nucleocapsid compared to anti-spike antibodies up to one year after SARS-CoV-2 infection. Diagn Microbiol Infect Dis. 2022;103(1):115659.

8. Koerber N, Priller A, Yazici S, Bauer T, Cheng CC, Mijocevic H, et al. Dynamics of spike-and nucleocapsid specific immunity during long-term follow-up and vaccination of SARS-CoV-2 convalescents. Nat Commun. 2022;13(1):153.

9. Erber J, Kappler V, Haller B, Mijocevic H, Galhoz A, Prazeres da Costa C, et al. Infection Control Measures and Prevalence of SARS-CoV-2 IgG among 4,554 University Hospital Employees, Munich, Germany. Emerg Infect Dis. 2022;28(3):572–81.

10. Hall V, Foulkes S, Insalata F, Kirwan P, Saei A, Atti A, et al. Protection against SARS-CoV-2 after Covid-19 Vaccination and Previous Infection. N Engl J Med. 2022;386(13):1207–20.

11. Matsumoto N, Sasaki A, Kadowaki T, Mitsuhashi T, Takao S, Yorifuji T. Longitudinal antibody dynamics after COVID-19 vaccine boosters based on prior infection status and booster doses. Sci Rep. 2024;14(1):4564.

12. Munro APS, Janani L, Cornelius V, Aley PK, Babbage G, Baxter D, et al. Safety and immunogenicity of seven COVID-19 vaccines as a third dose (booster) following two doses of ChAdOx1 nCov-19 or BNT162b2 in the UK (COV-BOOST): a blinded, multicentre, randomised, controlled, phase 2 trial. Lancet. 2021;398(10318):2258–76.

13. Walsh EE, Frenck RW, Jr., Falsey AR, Kitchin N, Absalon J, Gurtman A, et al. Safety and Immunogenicity of Two RNA-Based Covid-19 Vaccine Candidates. N Engl J Med. 2020;383(25):2439–5

14. Levin EG, Lustig Y, Cohen C, Fluss R, Indenbaum V, Amit S, et al. Waning Immune Humoral Response to BNT162b2 Covid-19 Vaccine over 6 Months. N Engl J Med. 2021;385(24):e84.

15. Bayart JL, Douxfils J, Gillot C, David C, Mullier F, Elsen M, et al. Waning of IgG, Total and Neutralizing Antibodies 6 Months Post-Vaccination with BNT162b2 in Healthcare Workers. Vaccines (Basel). 2021;9(10).

16. Regev-Yochay G, Gonen T, Gilboa M, Mandelboim M, Indenbaum V, Amit S, et al. Efficacy of a Fourth Dose of Covid-19 mRNA Vaccine against Omicron. N Engl J Med. 2022;386(14):1377–80.

17. Chalkias S, Harper C, Vrbicky K, Walsh SR, Essink B, Brosz A, et al. A Bivalent Omicron-Containing Booster Vaccine against Covid-19. N Engl J Med. 2022;387(14):1279–91.

